# Integrated deep visual and semantic attractor neural networks predict fMRI pattern-information along the ventral object processing pathway

**DOI:** 10.1101/302406

**Authors:** Barry J. Devereux, Alex Clarke, Lorraine K. Tyler

## Abstract

Recognising an object involves rapid visual processing and activation of semantic knowledge about the object, but how visual processing activates and interacts with semantic representations remains unclear. Cognitive neuroscience research has shown that while visual processing involves posterior regions along the ventral stream, object meaning involves more anterior regions, especially perirhinal cortex. Here we investigate visuo-semantic processing by combining a deep neural network model of vision with an attractor network model of semantics, such that visual information maps onto object meanings represented as activation patterns across features. In the combined model, concept activation is driven by visual input and co-occurrence of semantic features, consistent with neurocognitive accounts. We tested the model’s ability to explain fMRI data where participants named objects. Visual layers explained activation patterns in early visual cortex, whereas pattern-information in perirhinal cortex was best explained by later stages of the attractor network, when detailed semantic representations are activated. Posterior ventral temporal cortex was best explained by intermediate stages corresponding to initial semantic processing, when visual information has the greatest influence on the emerging semantic representation. These results provide proof of principle of how a mechanistic model of combined visuo-semantic processing can account for pattern-information in the ventral stream.

## Introduction

When we view an object, we understand the meaning of what we see. This complex process involves rapid analysis of the object’s visual properties, but object recognition involves more than visual processing alone. The activation of an object’s meaning is intrinsic in human object recognition. Visual processing leads to the automatic activation of an object’s conceptual knowledge^1^, but how this key transformation is achieved is not understood, since research on conceptual knowledge and vision largely progress independently of each other. Whilst research on vision focuses on the computational properties of occipital and temporo-occipital cortex, a wealth of neuroimaging and neuropsychological evidence has demonstrated that different kinds of semantic representations are activated along the ventral stream. In particular, whilst category structure (e.g. “tool”, “animal”) is represented in posterior fusiform cortex^2–6^, the anterior-medial temporal cortex (AMTC) is critical in activating amodal conceptual information about objects^4,7–9^. In particular, perirhinal cortex in the AMTC plays a critical role in the access of detailed, distinctive semantic information that is important in tasks which require unique identification of concepts, such as object naming (“hammer”, “tiger”).

To address the fundamental question of how the visual properties of an object elicit semantic information, we combine a computationally explicit model of vision with a computationally explicit model of semantics, to provide a proof-of-principle of one potential route by which visual properties intact with more abstract semantics. As a model of visual processing we use a deep convolutional neural network (DNN) which represents the state-of-the-art in machine vision research, and has remarkable power to capture visual information in images and to label objects accurately^10,11^. A DNN consists of a series of hierarchical layers, where the nodes of each layer correspond to filters that are sensitive to particular patterns in the preceding layer. Nodes in the earliest layers are sensitive to relatively local patterns of low-level properties of the stimulus (e.g. pixel values) whilst nodes at later layers are sensitive to higher-level complex visual features which exhibit invariance with respect to lower-level detail, such as position^12^. Although these models have been developed with engineering goals rather than neurocognitive plausibility in mind, recent neuroimaging studies have shown a remarkable correspondence between the layers of DNNs and activation patterns in the visual system^13–16^.

Importantly for our purposes, DNNs for vision do not include semantic knowledge about objects. They typically use an output layer consisting of object labels (e.g. BANANA, MOTORCYCLE), which do not capture information about the kinds of semantic commonality or semantic similarity which are an essential feature of human conceptual knowledge. An image of a tangerine and an image of a banana may be correctly labeled as TANGERINE and BANANA by the DNN, and will have orthogonal representations on the output label layer. Another object, such as a motorcycle, will also be orthogonal to the other two images, and so all three images will be maximally dissimilar to each other on the output layer (Figure 1). Training the network to activate orthogonal, maximally different representations on the output layer for different objects irrespective of their meaning solves the engineering goal of discriminating between objects, but this clearly fails to capture semantic similarity/dissimilarity between objects, such as the greater similarity between the concepts *banana* and *tangerine* compared to *motorcycle.* A visual DNN is a model of visual discrimination, rather than a model of the evocation of meaning that occurs during human object processing. Indeed, later layers of a visual DNN cannot account for semantic category distinctions frequently observed in IT cortex without an additional supervised training step where the model is explicitly trained to capture those categorical distinctions^14^. Thus, a visual DNN is an ideal model of the visual properties of objects, which we are able to exploit in order to determine the ways in which visual inputs activate semantic representations.

**Figure 1.**
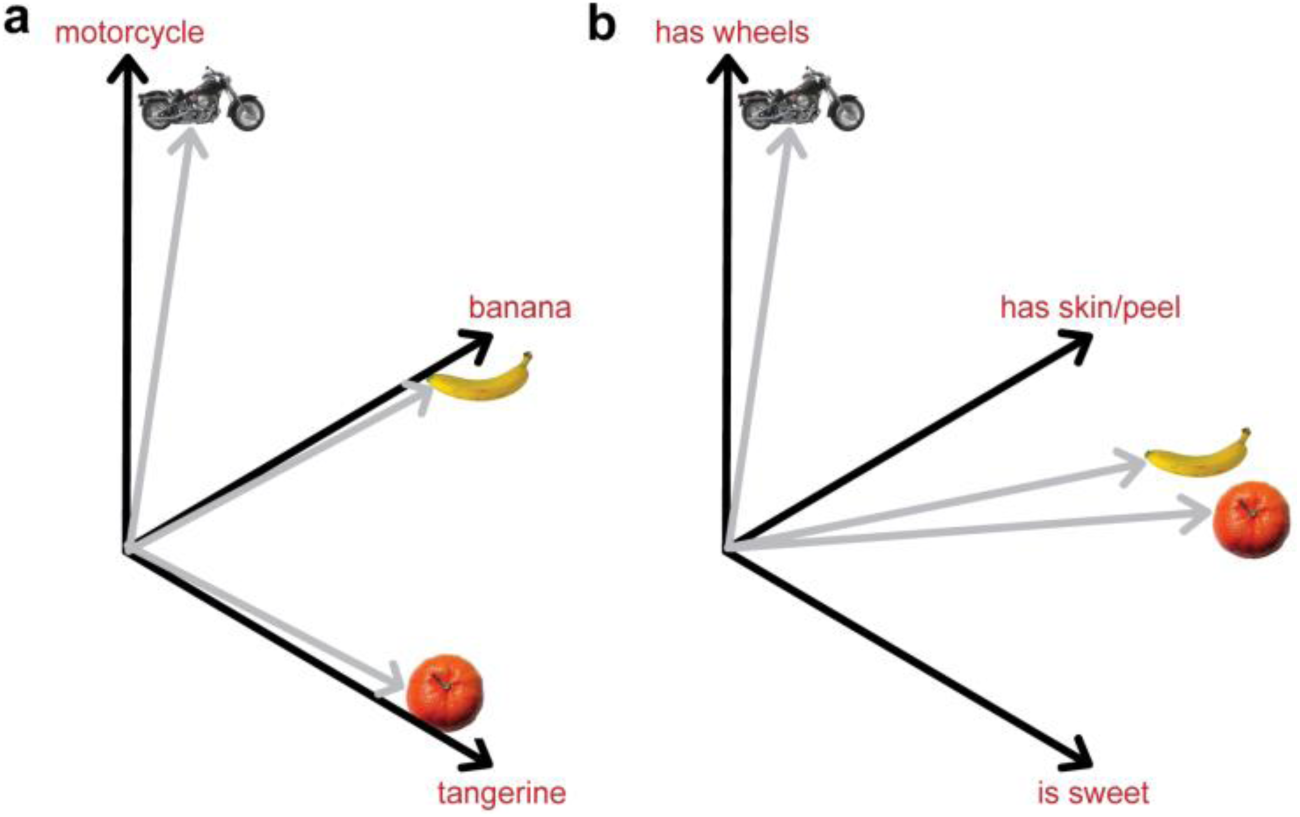
Representational spaces for object semantics. **(a)** The final layer of a visual DNN^10^ represents object in terms of 1,000 labels, and activations for different objects on the final layer can be visualized as points in the 1000-dimensional space spanned by the labels (for illustration, only 3 of the 1,000 dimensions are shown). All accurately-labeled objects will have orthogonal vectors in this space (grey lines) and so will be maximally dissimilar in this space, irrespective of the objects’ semantics. **(b)** A representation of objects in terms of their semantic feature vectors (e.g. using concept property norms). Each object is represented as a pattern of activation over semantic feature units (for illustration, only 3 dimensions are shown). Semantically similar objects, such as tangerine and banana will have high activations for shared semantic features and thus will be closer together in this space than to other semantically unrelated concepts (e.g. motorcycle).

To model the interaction between visual processing and meaning, we use a model of semantics that is based on a neurocognitively motivated distributed account of conceptual knowledge, where concepts are represented as patterns of activation across a network of semantic feature primitives^4,17–26^. In this framework a concept corresponds to a set of semantic features, and each semantic feature will activate for each concept that shares that feature. Distributed feature-based models naturally account for the semantic similarity between a pair of concepts in terms of the overlap of their features – two concepts that share many semantic features (e.g. *banana* and *tangerine*) will be closer in semantic space than concepts with little semantic overlap. These models also account for the differentiation between semantically similar concepts, with distinctive features that are true of only a few concepts (e.g. *has a mane*) helping to discriminate between a given concept and its close semantic neighbors (e.g. discriminating *lion* from other large cats)^1,3^. The distributed approach can be implemented as an attractor network model^27^, a dynamic recurrent neural network in which patterns corresponding to concept semantics gradually emerge through feature co-activation. To create a visuo-semantic model of object understanding, we combine a DNN model of vision with an attractor network such that the penultimate layer of the visual DNN serves as input to a dynamic conceptual system.

Critically, through the training of the attractor network, the model learns to associate high-level visual regularities found in later stages of the visual DNN with semantic information about objects. The combined computational model aims to describe how high-level visual information interacts with the statistical structure of concept features over the time-course of object processing, rather than to produce an accurate output labeling for arbitrary images. Using representational similarity analysis (RSA)^28^, we compare representations at each stage of the combined visuo-semantic model to fMRI data collected on a large set of familiar objects. We predict a gradient along the posterior-to-anterior axis of the ventral stream, with the low-level visual DNN fitting best with occipital activations, initial semantic activation in the semantic network best accounting for coarse-grained superordinate-category-level semantic activation in posterior ventral temporal regions, and later, detailed semantic activation in the network best accounting for anterior-medial cortex, including perirhinal cortex. Moreover, we predict that the gradual activation of meaning – in both brain and machine – will be sensitive to the statistical structure of concept features and the co-occurrence of semantic features with high-level visual properties.

## Materials and Methods

Using a DNN model of vision, we first obtained measures of activation for each layer of the network for a set of object images. The model activation patterns from the penultimate layer of the DNN were used as the input to train an attractor network model that maps between purely visual information and distributed semantic representations. We then tested the activation patterns of each layer of the visual DNN and each stage of the attractor network against the fMRI data using RSA

### fMRI data

We re-analysed fMRI data, originally reported in Clarke and Tyler^5^, collected from 16 participants (10 female, 6 male) who named aloud images of common objects. Participants gave informed consent and the study was carried out with the ethical approval of the Cambridge Research Ethics Committee. The data preprocessing and RSA analysis framework were the same as in Clarke and Tyler^5^ and therefore we only summarize the main aspects of experiment, data collection and pre-processing here. Each stimulus consisted of a photograph of an isolated object on a white background. A total of 145 object images were presented, 131 of which were from a set of six object categories (*animals*, *fruit*, *vegetables*, *tools*, *vehicles*, *musical instruments*). The remaining 14 objects did not belong to a clear semantic category and are not included in the analyses. All objects were presented once in each of six presentation blocks. The EPI images were not spatially normalized or smoothed in order to take advantage of high-spatial-frequency pattern information in the subsequent representational similarity analysis^28^. A general linear model was fit to each subject’s data separately (concatenating the data from the 6 presentation blocks) in order to obtain a single *t*-statistic image for each concept picture^28^. These *t*-maps, masked by a grey matter mask for each subject, were used in representational similarity analysis that compared the multivoxel patterns in the fMRI data to the patterns of activation on each neural network layer.

### Neural network modeling – visual DNN

In order to quantify the visual properties of objects, we used the deep convolutional neural network (DNN) model of Krievhesky et al.^10^, as implemented in the Caffe deep learning framework (`bvlc_reference_caffenet` implementation)^29^. This implementation has been trained on the ILSVRC12 classification dataset from ImageNet (1.2 million images). The network consists of 8 layers; five initial convolutional layers (*conv1*, …, *conv5*, with each convolutional layer consisting of between 43,264 and 253,440 nodes), followed by two fully-connected layers *fc6* & *fc7*; each consisting of 4096 nodes) and an output layer of 1,000 nodes corresponding to 1,000 object labels. The convolutional kernels learned in each convolutional layer correspond to filters receptive to particular kinds of visual input. In the first convolutional layer, the filters reflect low-level properties of stimuli, and include ones sensitive to edges of particular spatial frequency and orientation, as well as filters selective for particular colour patches and colour gradients^10,12^. Later DNN layers are sensitive to more complex visual information, such as the presence of specific visual objects or object parts (e.g. faces of dogs, legs of dogs, eyes of birds & reptiles)^12^, irrespective of spatial scale and orientation.

To obtain the activation values in this network for our set of images, we presented each of the object images to the pre-trained DNN and obtained the activation values for all nodes in each of the five convolutional layers and the final two fully connected layers (*fc6* and *fc7).* The images used were 627 images for 627 object concepts listed in a large property norm corpus^30^. All these images were in the same format as the stimulus images used in the fMRI experiment and included the 131 stimulus images as a subset.

### Neural network modeling – semantic attractor network

Estimates of semantic feature representations can be obtained from property norm data (e.g. the features of *banana* include *is a fruit*, *is yellow*, *grows on trees*, *has a skin/peel*, etc)^24,30^. Research on conceptual processing using detailed distributed feature-based semantic models^24,30^ has shown how semantic similarities between objects and the statistical structure of concept features can account for a range of behavioral and neuroimaging data^4,5,18,21,27,31–33^. The full set of features across all concepts serves as the basis for the semantic units of a flat attractor network, based on Cree et al.’s model of word meaning^27^. Each of the 2,469 nodes in the semantic layer corresponds to a semantic feature, with the meaning of each of the 627 objects represented as a combination of these features. We combine the Krievhesky et al DNN with this model of semantics based on distributed feature representations by removing the labelling layer from the visual model and replacing it with the attractor network. We trained the attractor network to learn to activate the correct binary pattern of semantic features for each image (1 if the feature is present in the concept, 0 otherwise). The network was trained using continuous recurrent back-propagation through time^34^ over 20 processing time-ticks. A node *n*’s external input (for a particular time-tick *t*) is calculated as a weighted sum of the current activation values of nodes which are connected to it:

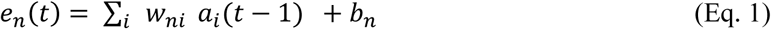

where *w_ni_* is the connection weight from node *i* to node *n*, *a_i_*(*t* – 1) is the activation of node *i* at time *t* – 1, and *b_n_* is the bias input to node *n*. The node’s total input, *x_n_* is calculated as a weighted combination of its external input and its total input at the previous timetick:

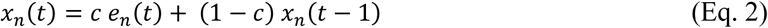

where *c* is a parameter that reflects how much the current total input is influenced by the total input at the previous timetick. Activation at time *t* is then calculated as a sigmoidal function of the total input:

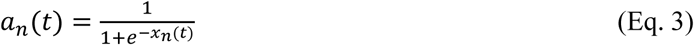

In the combined visuo-semantic (VS) model, input to the semantic system comes from the high-level visual representations of the final hidden layer of the DNN (i.e. layer *fc7*) for the set of 627 object images. To speed up training and reduce the number of parameters in the visual-to-semantic mapping, the 4096-dimensional data on layer *fc7* of the DNN was reduced to 60 dimensions through singular value decomposition. These 60 dimensions capture 58% of the variance of layer *fc7.* Representational dissimilarity matrices (RDMs) calculated on the full-dimensional *fc7* data and the SVD-reduced data are highly correlated (Spearman’s rho = 0.98), demonstrating that the reduction in dimensionality does not greatly affect the information content of the fc7 representations. The semantic attractor network component of the model thus has 60 input nodes (derived from *fc7*) mapping onto 2,496 nodes in the semantic network (*snet).* Connection weights were randomly initialised with values between 0.00-0.05 and activations for attractor layer nodes were initialised between 0.00-0.10. The attractor network was trained until at least 95% of the target semantic feature units in concepts reached an activation level of at least 0.70 (following Cree et al.^20,27^), which required 85 iterations of the image set. The simulations were built using the MikeNet neural network C library (version 8.02; http://www.cnbc.cmu.edu/∼mharm/research/tools/mikenet/). For further implementation details see Cree et al.^27^. Figure 2 depicts the full architecture of the combined VS model.

**Figure 2.**
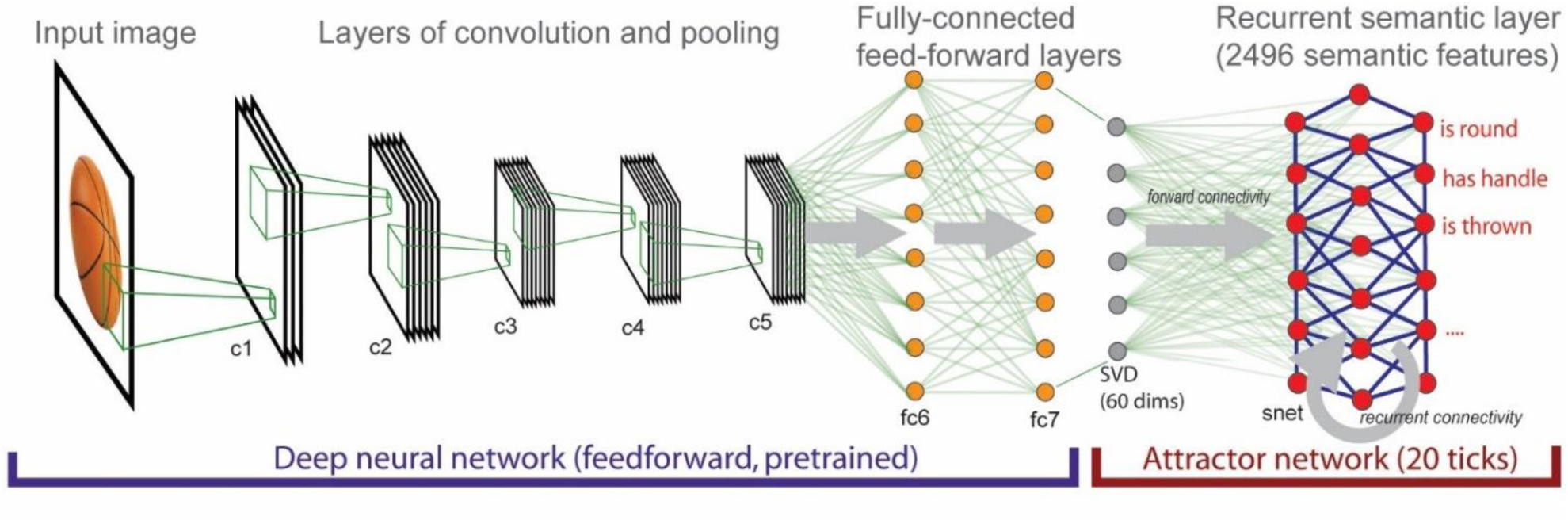
The combined visual DCNN + semantic attractor network architecture. The attractor network is a single fully recurrently connected layer (not all connections shown).

In the trained semantic model, semantic features activate gradually, over the course of 20 time-ticks. Activation of each feature is facilitated by feed-forward connectivity from the visual input layer and lateral, recurrent connectivity from other features in the semantic layer (i.e. correlational strength between features). The model therefore exploits statistical dependencies between high-level visual information and semantic features, and the speed with which individual semantic features activate depends both on their relationship to the visual input and their relationship to other semantic features. We expect early processing to be primarily driven by the visual input, with semantic features coding for shared, visual information (e.g. *is long*) activating more strongly than non-visual information (e.g. *is expensive*). At later stages of processing all semantic features of the object, irrespective of their type, will be maximally activated enabling the full, detailed meaning of the target concept.

### Representational similarity analysis

Representational similarity analysis (RSA)^28,35^ solves the problem of comparing very different kinds of multivariate data about a given set of items (e.g. voxel activation values for items in fMRI and node activation values for items in neural networks) by abstracting away from the underlying representational substrates (e.g. voxels & nodes) and instead representing information about items in terms representational dissimilarity matrices (RDMs) which capture items’ pairwise dissimilarities to each other (dissimilarity is typically calculated as 1-Pearson’s correlation between items’ multivariate patterns). Two RDMs can then be compared (e.g. by calculating a 2^nd^-order correlation) irrespective of the computational model or fMRI data which they are calculated over. The distributed nature of neural network data, where information is best characterized as patterns across units within a layer, makes RSA a natural analytical framework for relating such data to the brain^36^.

We calculated 26 RDMs for each stage of the VS model – one for each of the 7 layers of the visual DNN and for each of 19 processing time-ticks of the semantic attractor network model (the first time-tick of the attractor network model is randomly initialized and so is not included in the analysis). These RDMs were calculated over the same 131 images used in the fMRI experiment. Our combined visual+semantic architecture represents a hypothesis about how representations are transformed during object processing along the ventral stream, and the RDMs corresponding to different stages of the model can be conceptualized as a trajectory through RDM-space, moving from low-level visual detail to rich representations of conceptual semantics. RDMs from the model were tested against the fMRI data RDMs (gray-matter voxels) for each participant using the MRC-CBU RSA toolbox (revision 103) for MATLAB (http://www.mrc-cbu.cam.ac.uk/methods-and-resources/toolboxes). Spearman’s rho was used as the 2^nd^-order correlation measure for comparing brain and model RDMs. In order to more easily explore both the pattern of model fit for the different model stages within a region and the pattern of model fit across the cortex for an individual model, we used both standard region-of-interest (ROI) and searchlight mapping techniques, as implemented in the toolbox.

In the ROI analysis, we test the model RDMs against five fMRI RDMs for ROIs spanning the ventral object processing stream, namely bilateral early visual cortex (BA 17 & 18), left and right posterior ventral temporal cortex (pVTC), and left and right perirhinal cortex (PrC). The pVTC ROIs consisted of inferio-temporal, fusiform, temporo-occipital, lingual and parahippocampal cortex in the Harvard-Oxford atlas 70 to 20 mm posterior to the anterior commissure^37^, and the PrC ROI was defined using the probabilistic perirhinal map of Holdstock et al.^38^ thresholded at 2. Significance tests for the model RDM for each of the five ROIs were conducted using permutation of object condition labels (10,000 permutations)^39^ and then Bonferroni-corrected for five comparisons.

In the whole brain searchlight mapping analysis^28^ each of the 26 RDMs were tested across the brain (spherical searchlights, sphere radius of 7mm). For group random-effects analyses, the Spearman’s correlation maps for each participant were Fisher-transformed, normalized to standard MNI space, and spatially smoothed with a 6mm FWHM Gaussian kernel. To ensure optimal control of type I error, group-level random-effects analyses were conducted by permutation testing with the SnPM toolbox (http://go.warwick.ac.uk/tenichols/snpm)^40^. Variance smoothing of 6 mm FWHM and 5,000 permutations were used in these analyses. We report voxel-level false discovery rate (FDR) corrected *p*-values < 0.01 (one-sample pseudo-*t* statistic). For visualization, the volumetric statistically thresholded searchlight maps were mapped onto the PALS-B12 surface atlas in CARET version 5.6.

## Results

### Characteristics of the mapping between the DNN and semantic features

We investigated the learned relationship between high-level visual information in the DNN and features in the semantic model by examining the 148,140 connection weights learned by the attractor network between the 60 high-level visual input nodes and the 2,528 semantic feature nodes. First, we identified the individual semantic feature nodes with the largest connection weights connecting them to the visual input nodes (Figure 3a). Different nodes in the high-level visual input layer are sensitive to different visual qualities of images; for example, input node 1 has high values for visually round images, input node 5 has high values for green, leafy images, and so on. These high-level visual regularities can correspond to superordinate category information (e.g. many round things tend to be fruit or berries) although the information being represented is still purely visual in nature (e.g. the *helmet* image (image 12 for node 1; see Fig. 3a) is round and so has a high value on node 1, but is semantically unrelated to fruit/berries).

**Figure 3.**
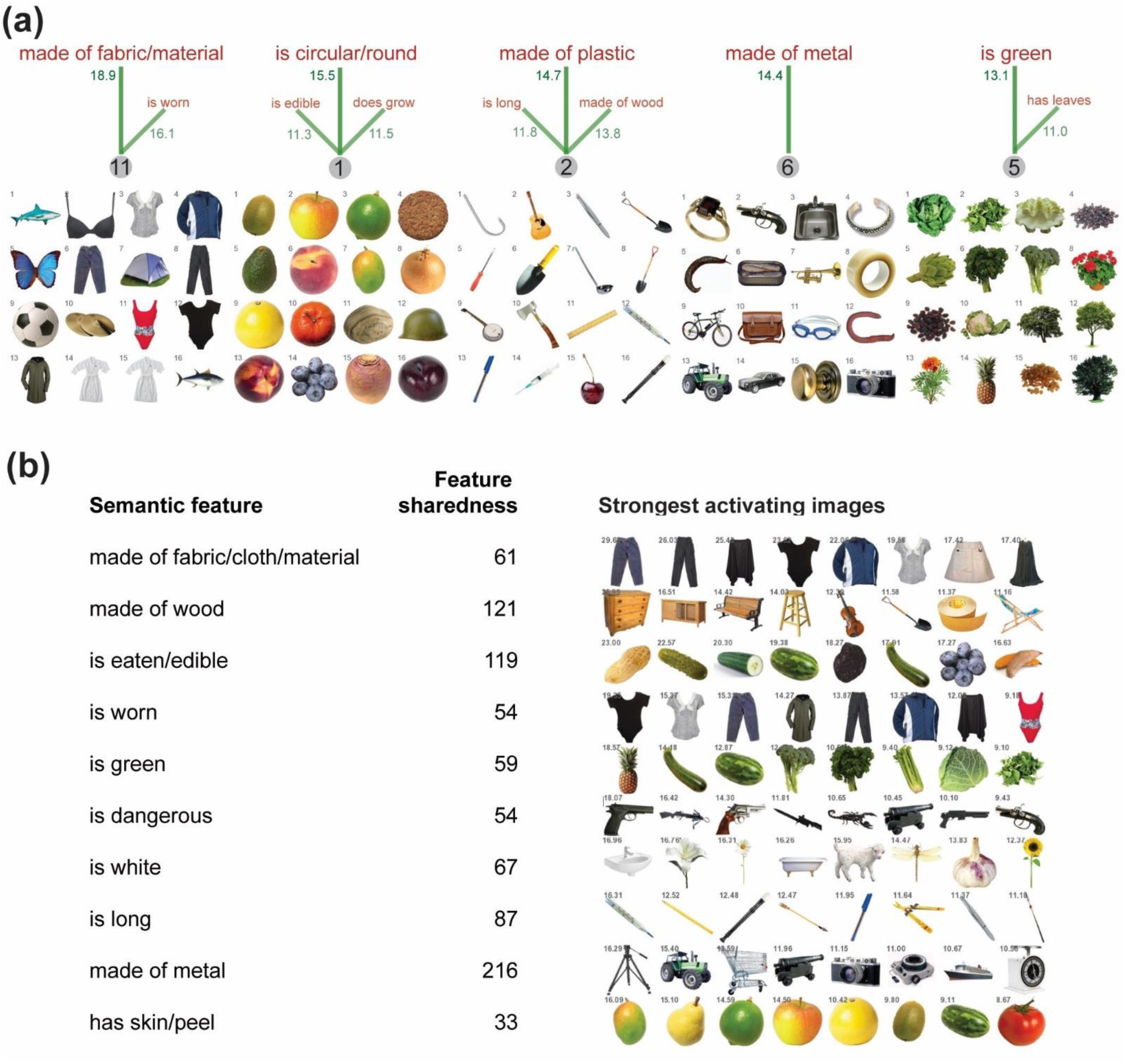
**(a)** Relationship between high-level visual information and semantic features in the attractor network model. The five semantic feature units with the highest learned connection weights to visual nodes are *made of fabric/cloth/material*, *is circular/round*, *made of plastic*, *made of metal*, and *is green.* For each semantic feature, the visual node (grey circle) with the strongest connection weight (green text) is shown. For each visual node, we display the 16 images with the highest values for that node. Also shown for each node are other semantic features with connection weights > 10. Different visual nodes reflect different high-level visual regularities in images which are mapped on to semantic features in meaningful ways (roundness, greenness, thinness, etc). **(b)** Semantic features which are most strongly activated by object images. For each semantic feature unit in the attractor model, we find the maximum input activation to that feature node from all visual input images. The 10 semantic features with the highest maximal input are shown, and the first image in each row is the features’ maximally activating image. The seven next most strongly activating images for each feature are also shown, as is the sharedness of each feature (i.e. the number of concepts the feature occurs in).

We also determined which images most strongly activate individual semantic features. For each image presented to the network, its net input to a semantic feature is a nonlinear weighted sum of its values across all 60 visual input units. Figure 3b presents the 10 semantic features which take the strongest external input from any of the input images, along with the images that give the strongest external input to each of those nodes. Certain images strongly activate certain semantic feature units in a manner which captures the statistical relationships between high-level visual information and particular semantic features. For example, semantic features such as *is long* and *is white*, which describe visual properties of objects, tend to be strongly activated by appropriate images (images of long things and white things, respectively). Other semantic features with strong visual input, such as *has a skin/peel* and *is dangerous*, also correspond to particular visual regularities (e.g. firearms with similar colours and physical form tend to strongly activate the *is dangerous* feature node). The semantic features with strong visual input also tend to be highly shared features which occur in many concepts. Highly shared features, such as *made of metal* tend to reflect information that is common to semantically similar concepts, whereas highly distinctive features, such as *has three wheels*, tends to be important for discriminating semantically similar concepts (e.g., distinguishing a tricycle from other similar vehicles). According to the conceptual structure account, shared feature information reflects coarse-grained semantics whereas distinctive information reflects later access to fine-grained semantic detail about objects^1,4,18^.

We also quantified the relationship between the visual input and the type of information conveyed by different semantic features. We calculated the *distinctiveness* of each semantic feature as 1/*N*, where *N* is the number of concepts in the norms that the feature occurs in^41^. Following previous research, we refer to highly distinctive features, occurring in just one or two concepts, as *distinguishing features*, and features occurring in three or more concepts as *shared features*^3,24,27^ For each concept, we obtained the activation level of each of its features at each of the 20 time ticks of the attractor network. We then averaged the activation values for shared features and for distinctive features across concepts. Although both shared and distinguishing features eventually become highly activated, shared semantic features are activated more rapidly at the earlier stages of semantic processing, reflecting the fact that these features are strongly activated from the visual input (Fig 4a). Similarly, we compared the semantic activation of different types of features using the classification of features into four semantic feature types that is given with the CSLB norms dataset^30^: visual (e.g. *is long*), non-visual perceptual (e.g. *is loud*), functional (e.g. *used for cutting*), and encyclopaedic (e.g. *associated with Halloween*). Visual semantic features are more rapidly activated from the input from the visual layer than other types of semantic features (Fig. 4b). Finally, we compared the activation of features with different levels of sharedness for different types (e.g. visual perceptual) (Fig. 4c). This clearly shows that high sharedness features are activated more rapidly compared to low sharedness and distinguishing features regardless of feature type.

**Figure 4.**
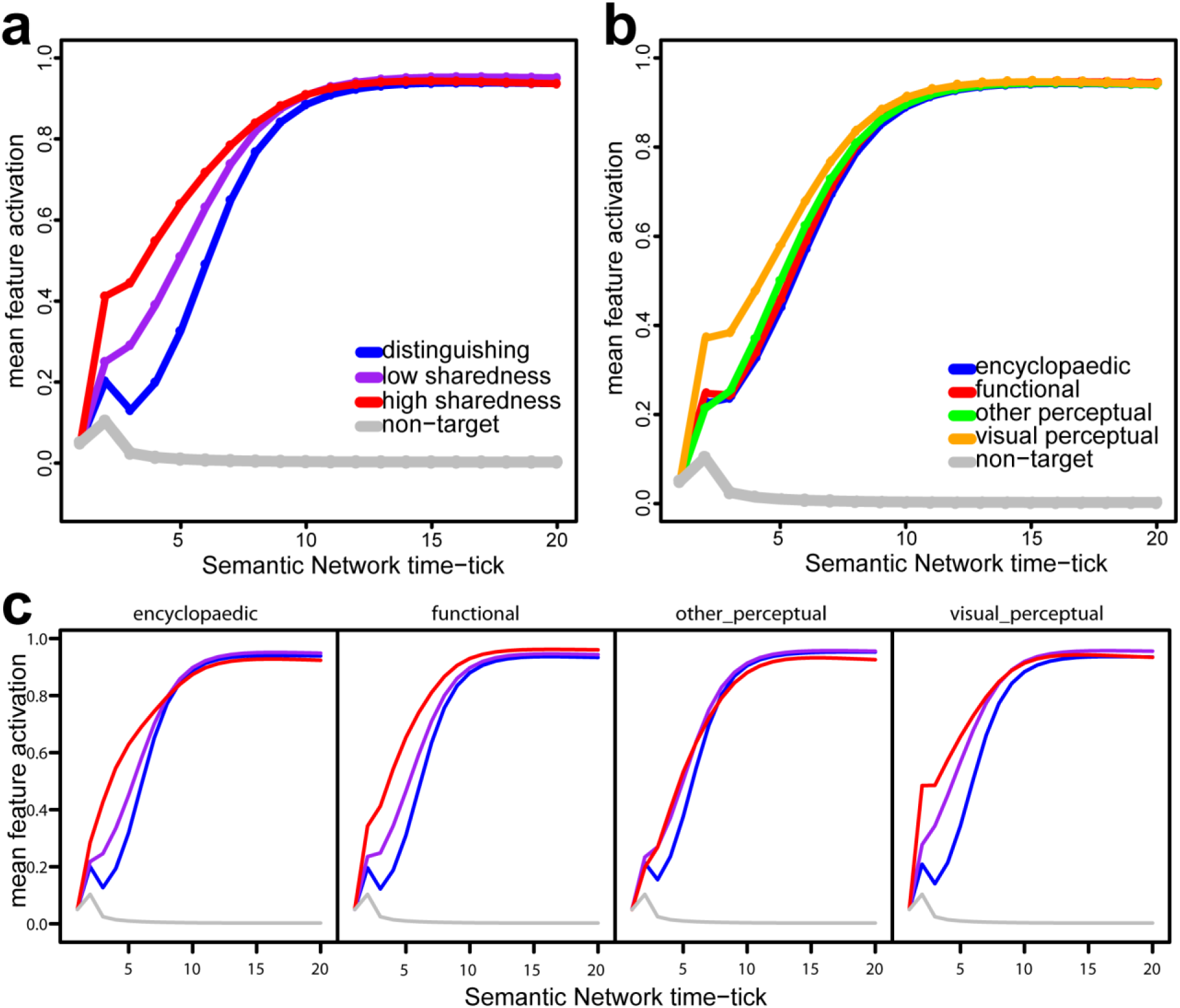
Activation levels of different types of semantic features. **(a)** Average activation level of distinguishing, low-sharedness, and high-sharedness target features (i.e. features that true of each concept) in the semantic part of the model, as a function of time-tick. Shared features were divided into high sharedness and low sharedness based on a median split. Highly shared semantic features activate more rapidly than distinguishing features. Average activation levels of non-target semantic features (i.e. features not true of the target concept; grey line) are also shown for comparison. (b) Average activation level of features of different types (visual-perceptual, non-visual perceptual, functional, & encyclopaedic). Semantic features classified as “visual perceptual” (e.g. *is round*, *has legs*) activate more rapidly than non-visual features (*e.g. is loud*, *grows on trees*, *used in baking*). (c) Average activation level of distinguishing, low-sharedness, and high-sharedness target features in the semantic network for features of different types. Across feature types, high sharedness features activate more rapidly than other features. Line colours match those in panel a.

These analyses of visual-to-semantic feature weights and semantic feature activation in the attractor network show that the model learns to associate high-level visual regularities found in the visual DNN with particular semantic properties. Moreover, the attractor network is sensitive to the conceptual structure statistics of images, with shared features activating more rapidly than distinctive features, and with distinctive (and non-visual) features relying more on lateral connections with other semantic features. The model therefore implements a general-to-specific account of object processing, where visual processing, and more coarse-grained semantic processing (relevant to broad superordinate categories) gradually gives way to more fine-grained concept-specific semantic representations^1,18^.

### fMRI analysis: Regions of Interest

Figure 5 presents the (Fischer-transformed) Spearman’s rho correlations between the neural RDMs for each ROI and the 26 model RDMs calculated for each stage of the full VS model. For the early visual cortex ROI (Fig. 5a) the best fitting stages of the model are for layers of the visual DNN. For both right and left pVTC, however, the best fitting model RDMs were from the early processing stages of the semantic attractor network (Fig. 5b,5c), when shared semantic features and visual semantic features are preferentially activated (see Fig. 4a). Finally, for both left and right perirhinal cortex, the best fitting model RDMs were from the last, and most semantically detailed, stages of semantic processing (Fig. 5d,5e).

**Figure 5.**
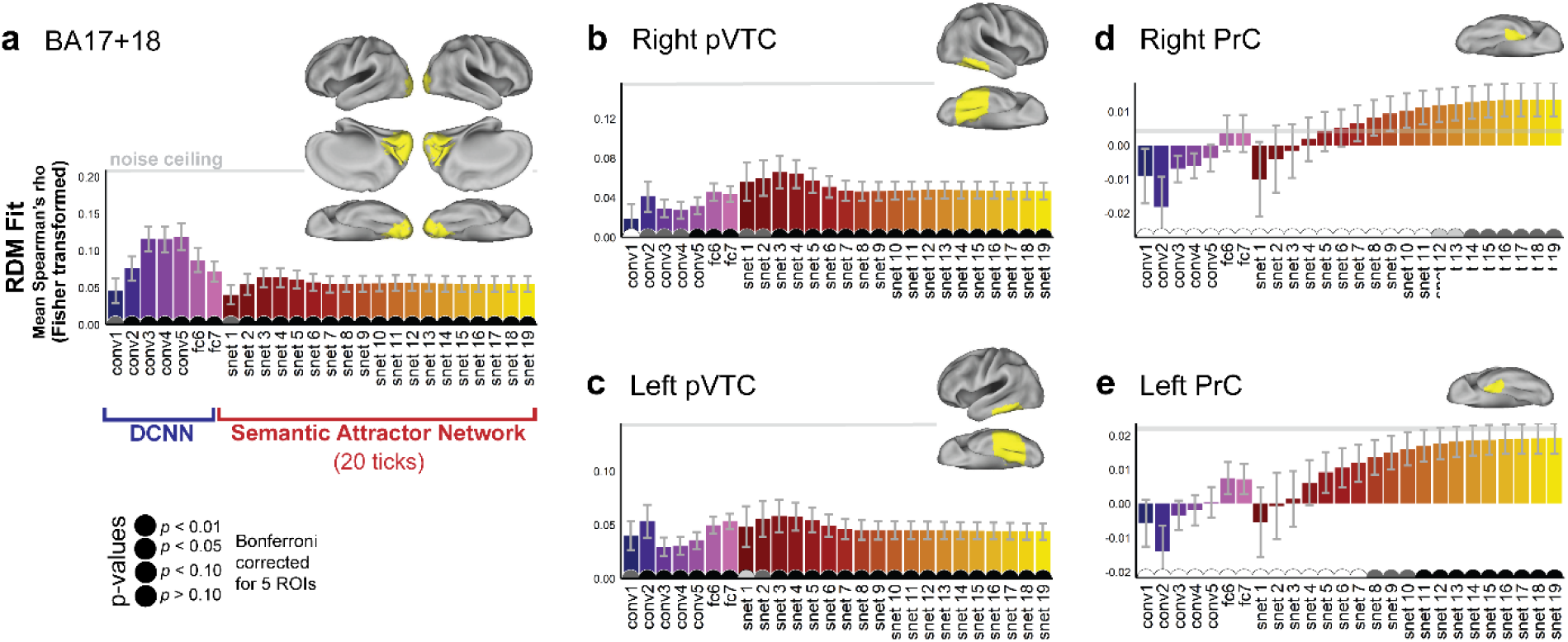
Region of Interest (ROI) results comparing representations at each stage of the VS model to patterns of activation along the ventral stream. ROIs are depicted as yellow regions on the cortical surface. Error-bars are SEM across subjects. Pips on the horizontal access depict *p*-values for tests of whether each individual Spearman’s correlation is greater than zero (permutation test, one-sided, Bonferroni-corrected for 5 ROIs). Grey lines indicate the lower bound of the noise ceiling.

We did not statistically compare each pair of RDM fits within each ROI, because the RDMs reflect a continuous trajectory through representational space, and so adjacent RDMs (e.g. the RDMs for fc6 and fc7 of the DNN, or ticks 18 and tick 19 of the attractor network) cannot be expected to differ significantly. Instead, we investigated whether the degree of RDM fit significantly varies as a function of model stage using a linear mixed effects analysis (i.e. we test whether RDM fit significantly increases or significantly decreases as a function of model stage). Our key prediction is that model stages differentially map onto different ROIs, such that earlier stages are a better fit to more posterior ROIs and later stages are a better fit to more anterior ROIs (i.e., an interaction between ROI and model stage on strength of model fit). In these analyses, model fit (i.e. Spearman’s rho correlations) were treated as a dependent variable, ROI (EVC, pVTC, PrC) as a categorical fixed effect, model stage (stages numbered 1 to 26) as an ordinal fixed effect, and subject as a random effect. Unlike multiple t-tests between pairs of RDM fits, this analysis takes into account the logical ordering of the model stages. Mixed effects analyses were implemented with the lmerTest package for R^42^. Firstly, we checked for hemispheric differences by including hemisphere as a fixed effect and testing for main effects of hemisphere and interactions between hemisphere and ROI and hemisphere and model stage (the bihemispheric ECV ROI was excluded from this analysis). No such interactions were found (all *p* > 0.05) and so we collapsed across hemisphere in subsequent analyses. We next tested for effects of region and model stage, and their interaction. We tested for non-linear relationships between model stage and RDM fit using restricted cubic splines (using the default quantiles of the rcs function in the R rms package). We iteratively fit models with increasing number of knots in the restricted cubic splines (i.e. increasingly complex non-linearities of model stage), stopping at the number of knots that gave the best fitting model (based on linear mixed effect model comparisons, as implemented in the lmerTest package).There was a highly significant main effect of ROI (*F* = 330.7, *p* < 0.001) reflecting the fact that overall model fit decreases as one moves from EVC to pVTC to PrC (which could be because of differences in the level of noise in the fMRI data across these ROIs). Consistent with the different patterns seen for each ROI in Fig. 5, there was a significant interaction of region and model stage (*F* = 22.1, *p* < 0.001). We explored the interaction effect by testing the simple effect of model stage in each region separately (Fig. 6). For the EVC ROI, there was a significant non-linear relationship between model stage and RDM fit, with model fit highest initially and decreasing for later model stages (*F* = 37.6, *p* < 0.001, 3 knots, Fig. 6 – blue curve). For the pVTC ROI there was a significant non-linear relationship between model stage and RDM fit, with model fit gradually increasing and peaking at the earliest semantic processing stages, and then decreasing slightly (*F* = 18.7, p < 0.001, 4 knots, Fig. 6 – green curve). Finally, for the PrC, there is a monotonically increasing relationship between model fit and model stage (*F* = 104.2, *p* < 0.001, 3 knots, Fig 6 red curve). The ROI results therefore reveal a posterior-to-anterior gradient along the ventral stream corresponding to the processing stages of the combined visual+semantic model, with more anterior regions corresponding to later processing stages of the model.

**Figure 6.**
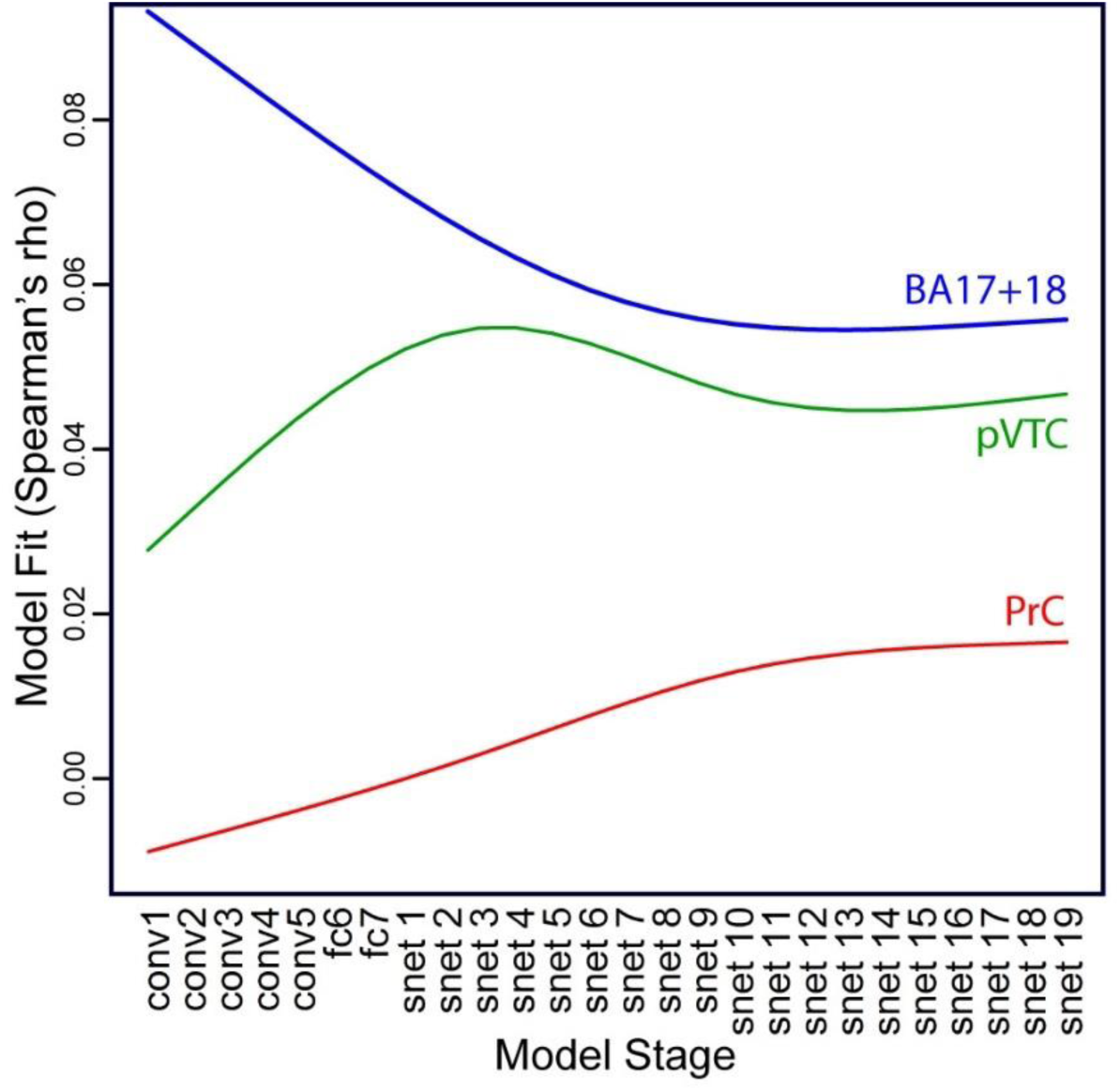
Relationship between model stage and model fit for the three ROIs: simple main effects of model stage for each ROI.

### fMRI analysis: Searchlight analysis

In order to investigate how each model stage relates to the fMRI data across the cortex, we also conducted searchlight RSA, correlating the RDM for each model stage to the neural searchlight RDMs across the whole brain. Figure 7a shows searchlight results for 4 of the 26 model stages (early and late visual stages, conv2 and fc7, and early and late semantic stages, at timeticks 3 (snet3) and timetick 19 (snet19); intermediate visual and semantic stages are intermediate to the depicted results). Consistent with the ROI results, significant effects do not reach the PrC until the final stages of semantic processing (Table 1). Figure 7b shows a composite map of the searchlight results for all 26 model RDMs. For each voxel where at least 1 of the 26 model RDMs was significant, we determined the best fitting model RDM (i.e. the model stage with the highest Spearman’s rho value). The results reveal a processing gradient along the posterior-to-anterior axis of the ventral stream, with visual stages of the model giving the best fit in occipital cortex, early stages of semantic processing (where general, shared semantic features and visual semantic features have stronger activation) giving the best fit to bilateral fusiform and finally later stages of semantic processing (including activation of distinctive feature information) giving the best fit to anteromedial temporal cortex including PrC (also see Sup Fig. 1).

**Figure 7.**
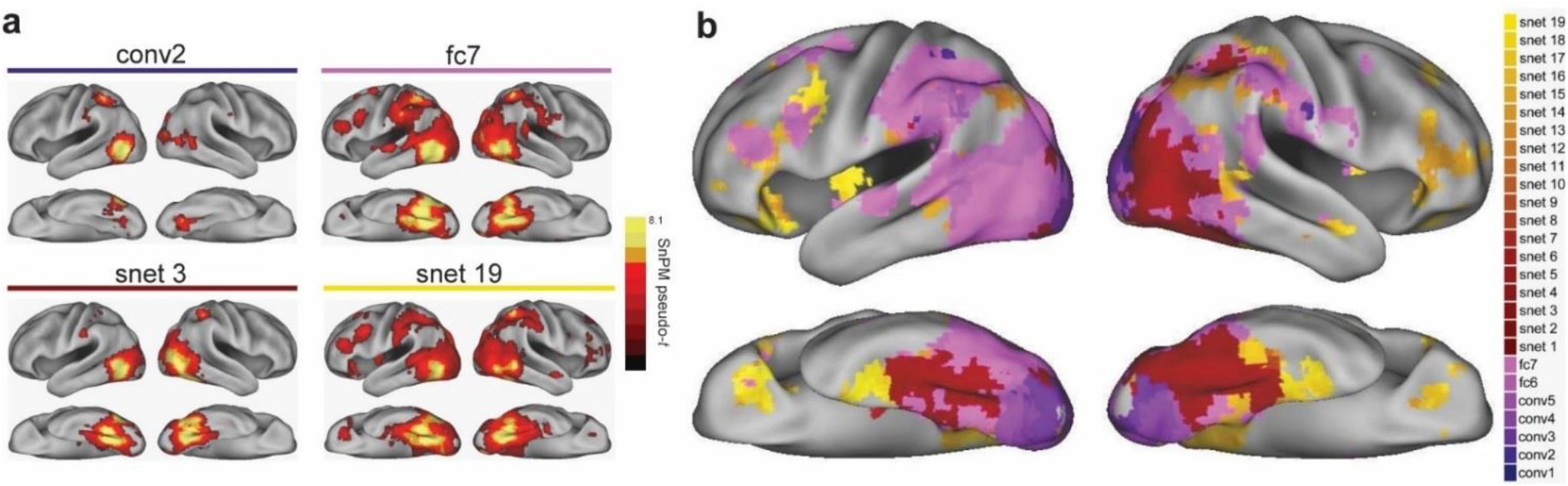
Searchlight results comparing representations at each stage of the VS model to representations across the cortex. **(a)** Individual RSA results for 4 stages of the visual+semantic model (early and late visual stages, conv2 and fc7; early and late semantic stages, snet3 and snet19). **(b)** Composite map of the searchlight results for all 26 model RDMs. Colors represent the best fitting model RDMs across the cortex. See also supplementary figure 1.

**Table 1.** RSA searchlight result for the 4 stages of the visual_semantic model in Figure 7.

| Regions | Cluster extent | Voxel-level p(FDR) | Pseudo-t | x | y | z |
| --- | --- | --- | --- | --- | --- | --- |
| **conv2** |  |  |  |  |  |  |
| L mid occip, sup occip | 821 | 0.0056 | 10.32 | -51 | -67 | -1 |
| R fusiform, mid occip, lingual | 510 | 0.0056 | 6.41 | 33 | -52 | -12 |
| L inf parietal | 113 | 0.0056 | 6.36 | -39 | -49 | 55 |
| **fc7** |  |  |  |  |  |  |
| L inf occip, L fusiform, R fusiform | 9589 | 0.0014 | 12.08 | -39 | -70 | -9 |
| L mid temporal | 114 | 0.0014 | 5.18 | -66 | -19 | -5 |
| L putamen, sup frontal, precentral | 118 | 0.0014 | 4.8 | -24 | -1 | 59 |
| L IFG (tri) | 119 | 0.0014 | 4.65 | -42 | 14 | 25 |
| L IFG tri, orb | 145 | 0.0014 | 4.26 | -48 | 32 | 14 |
| L ant cingulum, medial frontal | 190 | 0.0023 | 3.97 | 0 | 35 | 29 |
| **snet 3** |  |  |  |  |  |  |
| L inf occip, fusiform, R fusiform | 4251 | 0.0028 | 10.05 | -39 | -67 | -9 |
| R inf parietal | 102 | 0.0028 | 6.05 | 36 | -46 | 51 |
| **snet 19** |  |  |  |  |  |  |
| R fusiform, L inf occip, PRC | 10668 | 0.0014 | 10.07 | 27 | -64 | -9 |
| L IFG (orb), sup motor, precentral | 1430 | 0.0014 | 5.56 | -36 | 29 | -9 |
| R IFG orb, tri, mid frontal | 204 | 0.0084 | 4.02 | 33 | 32 | -9 |

MNI coordinates and significance levels shown for the peak voxel in each cluster. Anatomical labels are provided for up to 3 peak locations in each cluster. Abbreviations: mid = middle, occip = occipital, PRC = perirhinal cortex, inf = inferior, ant = anterior, sup = superior.

## Discussion

In the current study, we tested the degree to which a combined visual and semantic computational model could account for visual object representations. The model was successfully able to capture a posterior-to-anterior gradient of information from visual to semantic representations in the ventral stream, and additionally highlights a semantic gradient with early layers of the semantic network best represented in pVTC and late layers of the semantic network best represented in the PRC. This research shows a proof-of-principle for the combined computational model which offers one potential route by which visual properties interact with more abstract semantic information.

To investigate the transition between visual and semantic processes in object recognition we combined visual information from the layers of a DNN model of vision with a distributed, feature-based attractor network model of semantic processing, where the statistical dependencies between high-level visual information and semantic features are encoded in the connection weights between the high-level visual layer and a recurrent semantic system. The combined VS model therefore allows us to be explicit about the statistical regularities that facilitate the visuo-semantic mapping and makes quantitative predictions about the different stages of semantic activation. Activation of semantic features in this system is driven by both the high-level visual input as well as lateral recurrent connections with other semantic features. We found that shared semantic features and semantic features reflecting visual properties (e.g. “is long”) tended to activate more rapidly than more distinctive and less visual semantic features. This particular pattern of semantic activation demonstrates how visuo-semantic regularities and recurrent semantic processing can give rise to differences in how different kinds of semantic information activate over the processing stages of the model. In particular the model provides a computational implementation of a general-to-specific account of semantic processing, where coarse-grained, superordinate category-level information is initially activated from the visual input whilst fine-grained semantic information tends to emerge more slowly and is more reliant on the recurrent mutual co-activation of features within the semantic system^1,3,43^.

In complementary ROI and whole-brain searchlight RSA fMRI analyses, we found that the different processing stages of this model fit different areas of the ventral object processing stream. Firstly, visual stages of the model (i.e., the layers of the visual DNN) corresponded most closely to occipital cortex, consistent with other recent research testing DNN representations against neuroimaging data^13–15,44^. Going beyond this earlier work, we show that early processing within the semantic network – reflecting access to visuo-semantic information and coarse-grained semantics – provided the best fit to posterior ventral temporal cortex, bilaterally. Although we cannot rule out that a different model of high-level vision would not out-perform the early layers of the semantic network in the pVTC, our analysis does suggest more abstract semantic representations of objects related to coarse semantic information is present in addition to visual properties. This is consistent with previous neuroimaging findings showing that posterior fusiform cortex is particularly sensitive to shared-feature and superordinate-category level semantic representations^4,5,45^. However, in our model, superordinate-category-level semantics is not coded explicitly, and the superordinate-category-level representations in pVTC arise as a consequence of how visual representations interact with a dynamic and distributed feature-based semantic system that is sensitive to feature statistics (i.e. feature sharedness). Finally, PrC is best fit by the representations present in the final stages of semantic processing, where both distinctive and shared semantic information are maximally activated. This is consistent with the claim that PrC is critical to the integration of different kinds of semantic information in support of fine-grained object discrimination^4,5,46^. Overall, the stages of the model correspond well to a gradient in visuo-semantic processing across the ventral stream, but additionally highlights a conceptual gradient from the pVTC to the PRC.

While this research shows visual and semantic effects along the extent of the ventral stream, other work suggests a role for semantics in the angular gyrus, temporal pole and lateral anterior temporal cortex^8,47^. Our analyses did not yield significant effects in these regions which is likely due to the modality of the items. Semantic effects in the angular gyrus are more commonly associated with language^47^, and direct comparisons of the semantic effects between words and pictures show effects for words in the angular gyrus, but not pictures^45^. The temporal pole has been linked to the processing of amodal semantic information, in addition to the anterior temporal lobe as a whole^8^, although some research suggests different sub-regions within the ATL have particular sensitivities to different modalities^48,49^, with our PRC effects of semantics relating to the visual inputs.

Analysing neuroimaging data for a task by fitting computational models of the same task to the imaging data is a potentially powerful tool in cognitive neuroscience. Computational models are explicit about the mechanisms and information involved in the task and make specific quantitative predictions, whilst at the same time abstracting away from physiological detail that may be less relevant to a cognitive-level account^36,50^. However, an important consideration in this approach is the task that the model is trained to maximize performance on. The DNN models that have been of recent interest in vision neuroscience are typically optimized for the relatively narrow goal of visual discrimination performance in the ImageNet classification competition, which differs considerably from the goals of human object processing (e.g. understanding what is being seen, making inferences about how objects in a scene relate to each other, forming semantic associations, and so on, as well as visual identification). Information about the meaning of concrete concepts is elaborate and complex – our representation of the concept *apple* includes information about what *apples* look like, where they are found, how they are eaten, how they taste, what they are associated with, what they are a type of, what their subtypes are, and the contexts in which we are likely to encounter them. Depending on the demands of a given environmental context, such information must somehow be activated from the visual input. Even if semantic knowledge of objects is not logically required for visual discrimination (as evidenced by the human-level performance of DNNs in object classification), the activation of rich semantic representations may still be obligatory in human object processing. Indeed, patients with damage to the anteromedial temporal lobe, including PrC, show poorer naming performance associated with a loss of semantic knowledge, even when the visual system is intact, suggesting that semantic activation of specific conceptual representations is a pre-requisite for successful object naming in humans^6,46,51^. From a cognitive science perspective, visual DNN models may be overfit to the labeling task, and so can be trained to correctly identify objects without incorporating the kind of rich semantic system that is fundamental and obligatory in human object processing.

There are many ways to consider grounding a mechanistic semantic model in vision. Our architecture is constructed in two parts, using a pre-trained DNN and then an attractor network trained on the output of the DNN, and so there is no possibility for semantic representations to influence the representations in earlier layers, either through the backpropagation of error during training or through explicit feedback connections from the semantic network to previous visual layers. Although separate training of different model stages is in general not unusual (it is the basis of how deep belief networks are trained, for example) another approach would be to train the feed-forward visual components and the recurrent semantic components of the model together, so that all weights in both the visual and semantic parts of the network are learned simultaneously. Such a model would learn visual filters in the DNN layers that are useful for activating semantic knowledge about object concepts (which may not be the same as filters learned for labelling, in particular for higher levels). Alternative architectures might also explore direct connections from early visual layers to semantics, semantic-to-visual top-down connections, and lateral recurrent connections within visual layers (all of which would be neurocognitively plausible), whilst assessing the impact of using different visual models to map onto semantics. Furthermore, our model is trained to map high-level visual information about our specific stimuli images onto the corresponding semantic features (following the word meaning model of Cree et al.), and we have not tested the semantic patterns activated for previously unseen images (as is the case for most connectionist work on conceptual semantics). Our goal here is not to explore the infinite space of plausible model architectures, but rather to provide a proof-of-principle for how a distributional theory of concept meaning can be grounded in vision using a combination of state-of-the-art visual and semantic neural network models. The results show that our chosen model does this in a way that (a) predicts a general-to-specific account of semantic processing, and (b) explains the transformation of visual representations into semantic representations across the ventral stream.

In summary, we combined a deep convolutional neural network model of vision with a distributed attractor network model of semantics and tested the degree to which it captured object representations in the ventral stream. The model exploits statistical regularities between high-level visual information and semantic properties and makes predictions about semantic activation that are consistent with a general-to-specific account of object processing. RSA revealed that the different stages of the visuo-semantic model correspond convincingly to different stages of the ventral object processing stream, from early visual cortex to perirhinal cortex. By explicitly modelling the activation of rich semantic representations from the visual input, the model goes beyond identifying effects associated with visual models and semantic models separately, and instead shows how general and specific semantic representations are activated as a consequence of vision.

## Acknowledgements

This research was funded by an Advanced Investigator grant to LKT from the European Research Council under the European Community’s Seventh Framework Programme (FP7/2007-2013)/ ERC Grant agreement n° 249640, and an ERC Advanced Investigator grant to LKT under the Horizon 2020 Research and Innovation Programme (2014-2020 ERC Grant agreement no 669820).

## Author contributions

BJD, AC and LKT designed research; AC performed research and preprocessed data; BJD analyzed data and did the computational modelling; BJD, AC and LKT wrote the paper. All authors reviewed and edited the manuscript. The datasets generated analysed during the current study are available from the corresponding author on reasonable request.

## Supplementary figure

**Supp figure 1.**
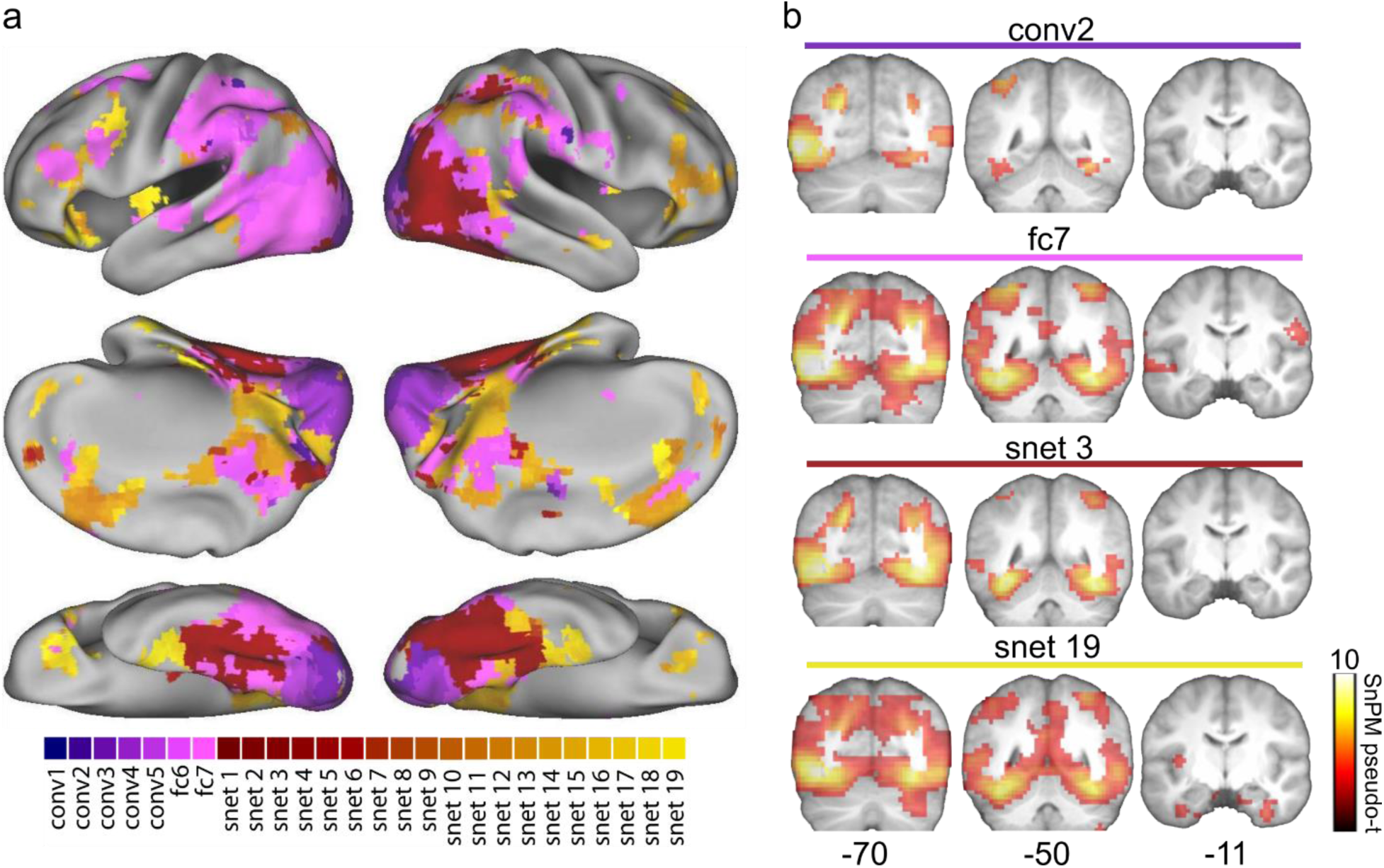
Searchlight results comparing representations at each stage of the VS model to representations across the cortex. **(a)** Composite map of the searchlight results, including a medial view, for all 26 model RDMs. Colors represent the best fitting model RDMs across the cortex. **(b)** Individual RSA results for 4 stages of the visual+semantic model (same as in Figure 7) at slices covering approximately early visual regions, posterior ventral temporal regions and anterior temporal regions.

